# TREM2 Alteration Increases AD Biomarkers and is Associated with Key Genes with 5xFAD Mice Model Analysis on MODEL-AD Database

**DOI:** 10.1101/2023.08.06.552135

**Authors:** Elena I. Budyak, Jihoon Kwon, Evan J Messenger, Surendra Maharjan, Juan J Koothur

## Abstract

Alzheimer Disease is a multifactorial disorder characterized by cognitive decline and memory loss. A key gene associated with AD is the TREM2 gene which has been identified as a risk factor for AD. Studies show that TREM2 functions in microglia regulation that controls the amount of AB proteins by the mechanisms of clearance and degradation. However, the exact mechanism of how the TREM2 variations like TREM2 KO and TREM2 R47H contribute to the progression of Alzheimer is still debated. Moreover, research into the levels of gene expression and measurements of biomarkers that contribute to progression of AD is very limited. In this paper, we conduct a comprehensive analysis of the biomarker concentration and gene regulatory behavior in TREM2 KO and TREM2 R47H mutated mice models from the MODEL AD database in order to gain understanding of how these variations contribute to the formation of biomarkers and contribute to AD progression.

Our research indicates a correlation between the mutated mice model and the different biomarker concentrations in the brain like insoluble AB40 and 42 proteins, soluble AB40 and 42 proteins, and NFL, which shows that TREM2 gene may be associated with multiple biomarkers. Moreover, we identified some key genes that were associated with the downregulation of the TREM2 gene with the TREM2 KO mice model gene analysis. Finally, we took the confocal images of the TREM2 KO mice model to analyze the effect that the lack of TREM2 extracellular receptor has on the neuritic dystrophy in the brain. Overall, we analyzed the biomarker concentration, gene regulatory activity, and the neuritic effects of the TREM2 KO and TREM2 R47H mutated variants of the TREM2 gene.

## Introduction

Alzheimer’s disease (AD) is the most common form of dementia, accounting for 60-80% of cases worldwide.^1^ In 2019, AD and other dementias were the seventh leading cause of death globally.^2^ AD causes progressive cognitive decline, culminating in difficulty performing daily activities and, eventually, death due to complications such as aspiration pneumonia.^1^ Alzheimer’s disease is characterized by the accumulation of beta-amyloid (Aβ) and tau proteins in neuritic plaques and neurofibrillary tangles, respectively, resulting in neuron damage and death. In 2013, it was found that the R47H variant in the triggering receptor expressed on myeloid cells 2 (TREM2) gene was associated with an increased risk for AD.^3,4,5^ Before this, much of the research on TREM2 was on mutations leading to a rare form of autosomal recessive early-onset dementia known as Nasu–Hakola disease (NHD), which presents with bone cysts and fractures.^3^ Due to this discovery, TREM2 function and variants, especially R47H, have increasingly become the focus of research on the progression and management of AD. However, much is still unknown, including TREM2-R47H’s mechanism of exacerbating AD pathogenesis and applicability in diagnostics. Therefore, the goal of this paper is to establish a correlation between TREM2 mutations and the concentrations of the AD-associated biomarkers: insoluble Aβ40, insoluble Aβ42, soluble Aβ40, soluble Aβ42, and neurofilament light chain (NFL), which could prove helpful in diagnosis and predicting prognosis.

### Amyloid Pathology

#### Amyloid Hypothesis and Aβ Generation

Beta-amyloid proteins, along with tau proteins, form the basis of the amyloid hypothesis, which is that the accumulation of misfolded extracellular Aβ in neuritic plaques and tau in neurofibrillary tangles results in neuronal damage, leading to confusion, memory loss, and cognitive decline characteristic of AD.^6^ Aβ peptides are 37 to 49 amino acids long and are produced from the processing of amyloid precursor protein (APP).^6^ APP is metabolized by an amyloidogenic and a nonamyloidogenic pathway.^6^ The nonamyloidogenic mechanism involves APP being cleaved by α-secretase, releasing a sAPPα molecule from the cell surface and leaving a membrane-bound 83-amino-acid C terminal fragment (C83).^6^ The α-CTF is then cleaved by γ-secretase to produce P3 and leave an APP intracellular domain (AICD). The amyloidogenic pathway involves β-secretase cleaving APP, sAPPβ release, and C99 formation. C99 is cleaved by γ-secretase to yield Aβ peptides of various lengths, the forms most relevant to AD are the 40- and 42-amino-acid Aβ40 and Aβ42. AICD from C99 origin can translocate to the nucleus and possibly affect gene expression, including that of apoptotic genes.^6^ In AD amyloidosis, monomeric Aβ that is not degraded or removed from cerebrospinal fluid (CSF) bind together to form soluble oligomers and insoluble fibrils and plaques.^7^ While the original amyloid hypothesis is that the deposition of Aβ into plaques leads to neurotoxicity and tau proteins forming toxic neurofibrillary tangles, this hypothesis has been challenged in multiple ways.^7^ For instance, monomeric Aβ was previously regarded as a functionless product of APP cleavage, but many studies indicate that it serves a number of physiologic roles.^8^

### Monomeric Aβ Physiology

While monomeric Aβ was previously regarded as a functionless product of APP cleavage, some studies indicate that it serves a number of physiologic roles.^8^ Many of these studies and hypotheses on monomeric Aβ suggest that it behaves in a hormetic manner; it is pathological in high concentrations and beneficial at low concentrations.^8, 9^ According to a study using autaptic cultures of hippocampal neurons, it was found that Aβ monomers regulate synaptic formation.^10^ This suggests that Aβ monomers might limit excitatory synaptic transmission by limiting the number of synapses, thus reducing cellular energy demand and excitotoxicity.^10^ Other studies find a hormetic correlation between the Aβ concentration and synaptic activity and demonstrated that long-term potentiation (LTP) is impaired by infusions of high concentrations of Aβ (likely to aggregate into oligomers), anti-Aβ antibodies, and other anti-Aβ peptides, while low concentrations of Aβ led to normal LTP.^8^ Long-term potentiation is a process by which post-synaptic neurons become more prone to stimulation by pre-synaptic neurons and is associated with memory formation. Two studies taken together indicated that Aβ12-28 in nanomolar concentrations enhanced memory formation and learning while diminishing those in micromolar concentrations when testing mice with foot-shock avoidance mazes.^8^

Some studies have indicated that monomeric Aβ functions as an antimicrobial agent (AMP).^8^ The “Antimicrobial Protection Hypothesis,” building on the prior “Bioflocculant Hypothesis,” suggests that Aβ binds to toxins and pathogens and aggregates into plaques to assist in the phagocytic removal of these substances and organisms and that dysregulation of this mechanism leads to neuroinflammation and degenerative plaque formation.^8^ These hypotheses are supported by structural similarities between antimicrobial peptides and Aβ and their similar function of aggregating in response to infections.^8^ In vitro experiments have demonstrated synthetic and cell-derived Aβ binding to and agglutinating various bacteria, *Candida albicans* fungi, influenza, and HSV-1.^8^ Furthermore, studies with 5XFAD mice expressing human Aβ have shown greater survival rates than wild-type mice with *Salmonella Typhimurium* and HSV-1 infusions. However, in one of the HSV-1 studies, cell-derived Aβ (containing oligomeric Aβ) was more effective at lower concentrations than synthetic monomeric-only Aβ in inhibiting the virus, potentially indicating that oligomeric Aβ can serve as an effective antimicrobial agent in normal physiological conditions. In addition to exhibiting antimicrobial properties during in vitro and in vivo experiments, Aβ has indirectly been shown to have those properties because patients in clinical trials of β-secretase and γ-secretase inhibitors have experienced higher infection rates.^8^

Other proposed functions of monomeric Aβ include improved recovery from traumatic brain injuries (TBIs), angiogenic processes, and inhibiting the proliferation of cancer cells. The TBI recovery hypothesis is supported by reports of elevated Aβ in TBI survivors and can even remain elevated in axons without resulting in plaque formation in the long term.^8^ However, conflicting evidence exists in studies with BACE1 (β-secretase enzyme) KO mouse models.^8^ One study indicates that the BACE1 knockout improved behavioral deficits and cell death, while another suggests that it resulted in reduced motor performance compared to controls.^8^ The latter study’s authors attributed this to the age difference (the former study used aged mice, which would be more prone to poor TBI recovery, with or without BACE1 KO and the corresponding absence or presence of Aβ). The angiogenesis hypothesis is supported by studies showing Aβ at nanomolar concentrations inducing endothelial cell proliferation and angiogenesis, while micromolar concentrations killed endothelial cells. Other studies found that zebrafish with APP deficiency could have vascular branching abnormalities alleviated with human monomeric Aβ infusions. The effects of Aβ on cancer have been demonstrated by studies showing that AD patients have lower cancer rates than non-AD, elderly individuals.^8^ Additionally, infusions of gliomas into transgenic Aβ overexpressing mice resulted in more tumor reduction than non-transgenic mice.^8^

While the results of these studies indicate that monomeric Aβ certainly has beneficial physiologic roles, some of these studies have significant design flaws. For example, oligomeric Aβ solutions might be used that actually contain more monomeric Aβ than oligomeric, according to transmission electron microscopy. Monomeric Aβ tends to dimerize and oligomerize, making determining precise concentrations challenging. Some studies might fail to consider pre existing endogenous Aβ while assessing the effect of Aβ infusions.^8^ Synthetic Aβ might not accurately represent in vivo Aβ effects as it might contain different isoforms. BACE1 inhibition might not prove the roles of Aβ as its other substrates, whose metabolism would also be impaired, might be responsible for the effects attributed to Aβ.^8^ More studies must be conducted with improved experimental methods, especially concerning Aβ preparation, to determine the functions of monomeric Aβ, its mechanisms of action, and the causality of conflicting effects.

#### Insoluble Aβ is not Responsible for AD Pathogenesis

In addition to the functionality of monomeric Aβ, the amyloid hypothesis has also been challenged regarding the role of insoluble Aβ. It was initially believed that Aβ plaques lead to neurotoxicity and the symptoms of AD.^6, 7^ Experiments have indeed shown correlations that would suggest this. Jun Wang and associates from the University of Pennsylvania’s Center for Neurodegenerative Disease Research and the Mayo Clinic used a sandwich enzyme-linked immunosorbent assay (ELISA) to detect levels of soluble, insoluble, and total Aβ, Aβ1-40, and Aβ1-42 in samples from age-matched normal, AD, and pathologic aging (no cognitive deficits but amyloid pathology present) human brains.^11^ They found that all Aβ1-40 and Aβ1-42 biomarkers were highest in AD brains, intermediate in pathologic aging, and lowest in normal brains. Their study also found that while soluble Aβ40 and soluble Aβ42 comprised most of the Aβ pool in normal brains (50 and 23%, respectively), it comprised the lowest in AD (2.7 and 0.7%, respectively) and intermediate in pathologic aging (8 and 0.8%, respectively). They concluded that AD pathogenesis is likely caused by a shift in the Aβ from soluble to insoluble forms and that pathologic aging was a transition state from normal aging to AD.

However, later studies would demonstrate that this shift from soluble to insoluble Aβ is not the cause of AD, or at least the amyloid plaques are not the principal cause. For instance, the neuroscientists Engler et al. from the Karolinska Institute and other institutions in Stockholm have found “Relatively stable PIB retention after 2 years of follow-up in patients with mild Alzheimer’s disease.”^12^ Pittsburgh Compound-B, or PIB, is an imaging agent used in positron emission tomography that binds to Aβ peptides (but binds particularly well with Aβ fibrils in plaques).^13^ The researchers conclude that the stable PIB retention indicates Aβ deposition leveling off at an early stage in AD and that it “may precede a decline in rCMRGlc [regional cerebral metabolic rate for glucose] and cognition.”^12^ This is relevant because it means that amyloid plaque formation does not progress with the decline in cognition seen in AD, likely meaning that the amyloid plaques are not the causative agent behind cognitive and memory declines. Other studies and analyses further confirm this, such as those which show that some individuals with high amyloid plaque burden have no cognitive deficits.^14,15,16^ In fact, if estimates on the amyloid burden in cognitively normal people from autopsy studies are correct, up to 50% of people 65 years or older have amyloid deposition, and half of them do not suffer from cognitive impairment.^16^ Studies like these have led to a consensus among neuroscientists that amyloid plaques are not the primary cause of AD.

On the contrary, researchers remain conflicted on the causation of AD. Some remain in support of aspects of the amyloid hypothesis.^15^ Others have abandoned it in favor of other theories, such as the Tau and inflammatory hypotheses.^15^ Concerning the amyloid hypothesis, there are two major ways that Aβ could still be the root cause of AD. One of these is that Aβ might initiate mechanisms that, once started, can propagate into a pathologic state independently of Aβ.^17^ Indirect evidence could exist in a human Aβ immunization trial (AN-1792).^17^ Eighty patients who met the NINCDS-ADRDA “criteria for probable Alzheimer’s disease with mild to moderate dementia (14–26 points on the mini-mental state examination)” entered a clinical trial for treatment.^18^ The AN-1792 immunization contains synthetic full-length Aβ peptides and is intended to initiate an immune response that should clear amyloid deposits.^19^ In the patients who received the immunization, amyloid deposition was lower than expected for the patients’ ages post-mortem, though the patients still experienced end-stage dementia.^17^ However, this evidence must be analyzed with caution as the patients’ pre-treatment amyloid loads were not able to be determined, meaning that the treatment might not have reduced amyloid load as much as believed, if at all if the patients began their AD prognosis later than would be expected based on their mini-mental state examination.^17, 18^ The hypothesis of Aβ initiating a mechanism that can propagate independently of the state of amyloid aggregation can answer why amyloid removal supposedly does not affect whether patients develop end-stage dementia. It can also explain why Engler et al. found little progress in amyloid deposition despite a cognitive decline over the same two-year period. However, it cannot address why amyloid deposition is present in some cognitively normal individuals (any Aβ-initiated neurodegenerative process tied to the amyloid deposition cascade should be activated if deposits have formed). Neither can the other prominent remnant of the amyloid hypothesis that comprises the mainstream view on AD pathogenesis.

Many researchers believe that soluble Aβ oligomers are the primary toxic substance and that plaques might be a sink for containing the oligomers in a less harmful form.^15, 20^ There is evidence that would support the hypothesis of oligomeric toxicity. For example, the neuroscientists and pathologists Shankar et al. from Harvard University and notable medical and research institutions in Ireland studied the effects of soluble human Aβ infusions on rat brains.^21^ They found that oligomers, specifically dimers, diminished LTP, increased long-term depression (LTD), and decreased the hippocampal density of dendritic spines. Long-term depression is the counterpart to long-term potentiation and involves reduced sensitivity of postsynaptic neurons to presynaptic signaling at underused synapses. LTD has been shown to inactivate aversive associative memories, while LTP was shown to form and re-activate those memories.^22^ However, it has also been demonstrated that LTD is necessary to consolidate spatial memory.^23^ Therefore, it is possible that increased LTD, reduced LTP, and the dendritic spine density reduction (likely impairs signaling) in the hippocampus are responsible for Shankar et al.’s observation that soluble Aβ dimers disrupted the memory of learned behavior in normal rats.^21^ Additionally, they found that the Aβ plaques did not diminish LTP unless disassembled into dimers with formic acid, indicating that plaques function as a nontoxic sink for toxic amyloid oligomers.^21^ While this study and others like it would suggest that oligomeric Aβ species such as dimers could be the neurotoxic species responsible for AD, as previously stated, they cannot explain how cognitively normal individuals can have significant amyloid deposition, which would have to be preceded or accompanied by increased levels of Aβ oligomers that would cause neuron damage.

Other studies, such as by neuroscientists Kim et al. from the Mayo Clinic and the University of Florida, indicate that Aβ might not be as toxic as once thought.^24^ The researchers used transgenic mice with BRI2 genes (a gene implicated in familial British and Danish dementias) where an Aβ1-40 or Aβ1-42 sequence replaced the Abri peptide sequence at the C-terminus. This transgenic model would overexpress the Aβ1-40 or Aβ1-42 or both peptide sequences which attached to the BRI2 (those peptide sequences would be cleaved off by proprotein convertases which naturally liberate Abri from BRI2).^24^

The researchers found that despite the overexpression of Aβ1-40 or Aβ1-42 or both, the mice did not suffer cognitive impairments pre- or post-plaque formation. They believe that this implies that Aβ alone is not the pathogenic substance that causes neurodegeneration.^24^ Kim et al. suggest that the cognitive deficits that were observed in APP overexpression models occur through molecular mechanisms that BRI2-Aβ cannot replicate, such as high levels of APP processing derivatives, which might be the actual cause for toxicity, being bound to the Aβ oligomers that are falsely believed to be toxic.^24^ Additionally, they state that models—such as Shankar et al.—which administer synthetic or natural Aβ into rodent brains could demonstrate memory deficits but not be in conflict with their study because infusions cause acute exposure rather than chronic aggregation of Aβ, and hence, do not reflect the normal progression of amyloid pathology.^24^ In summary, amyloid oligomerization might not be the initiating factor for cognitive impairment in AD patients, but oligomers could still play a role as aggregation points for other APP-derived toxins.

As previously stated, the paradigm on the cause of AD has significantly shifted from the original perspective on beta-amyloid and its mechanisms of pathogenesis. Monomeric Aβ is now believed to engage in beneficial functions, such as synaptic regulation, LTP, immunity, TBI recovery, angiogenesis, and tumor suppression, under normal physiological conditions, often in a hormetic manner. Insoluble Aβ is no longer considered the pathogenic agent of AD, and even oligomeric Aβ’s role is debatable. Due to significant evidence for other theories, we refer you to the reviews by Kametani and Hasegawa on the tau hypothesis and Morris et al. on the various hypotheses on AD.^7, 15^ While the oligomeric amyloid hypothesis’s validity is far from confirmed, it is essential to continue research into Aβ and its interactions with other molecules to determine the mechanisms that lead to Aβ being a feature of Alzheimer’s and potentially discover new therapeutic targets associated with Aβ oligomers. Since Aβ remains helpful in clinical practice as a biomarker for AD, it is important to understand how genes associated with familial AD, such as TREM2-R47H, affect its concentrations in vivo.

### TREM2

#### TREM2, Ligand Interactions, and R47H

The TREM family of genes codes for extracellular surface receptors on myeloid cells, such as microglia.^5^ These receptors are composed of an extracellular immunoglobulin variable-type domain situated on a stalk, a short cytoplasmic domain, and a transmembrane domain containing a positively charged residue that can bind to a negatively charged residue in a transmembrane adapter.^5^ The TREM2 AD risk variant, R47H, is an Arg → His substitution at position 47, located in the extracellular ligand-binding domain.

TREM2 has been shown to play a critical role in various microglial functions. An in vitro study on NHD found that TREM2 knockdown caused inhibition of microglial phagocytosis of apoptotic neurons and increased transcription of tumor necrosis factor α (TNF-α) and nitric oxide synthase-2.^25^

TNF-α is a pro-inflammatory cytokine, and nitric oxide performs both pro- and anti-inflammatory roles in the body, in addition to various other functions such as mediating apoptosis and serving as a neurotransmitter.^26,27,28^ Inflammation is a significant AD feature, and the accompanying oxidative stress can affect neuronal function. Specifically, nitric oxide can react with superoxide to produce peroxynitrite, which can go on to damage lipids, proteins, and nucleic acids. An example of this is the nitration of tyrosine’s phenolic rings, leading to abnormal proteins implicated in AD.^28^ The NHD study that found that TREM2 knockdown inhibits phagocytosis and increases damaging neuroinflammation also found that TREM2 expression increased chemokine receptor expression, cell migration, phagocytosis of apoptotic neurons, and diminished pro-inflammatory responses.^25^ The results of this study indicate that TREM2 plays a critical role in maintaining homeostasis in the brain by preventing inflammation while supporting microglial phagocytosis and migration.

TREM2 is a receptor that binds to various ligands, many of which are lipids. It has been reported to interact with phospholipids and sulfatides (sulfatides comprise part of myelin sheaths; likely how microglia respond to apoptotic neurons), lipoproteins, such as high-density lipoprotein, low-density lipoprotein, and apolipoprotein E (APOE), and bacterial lipopolysaccharide (LPS).^5, 29^ TREM2 has also been shown to bind to Aβ proteins, serving as a mechanism by which microglia can cluster and eliminate amyloid oligomers and deposits. The R47H and R62H (another AD risk variant) mutations result in a reduced ability for TREM2 to bind with oligomeric Aβ.^30^ While the exact structural mechanisms by which TREM2 binds to lipids, Aβ, and other molecules and how R47H affects these mechanisms are unknown, there are several hypotheses. According to one study, risk variants such as R47H do not significantly affect the binding affinity of TREM2 with phospholipids, but intracellular signaling is diminished.^31^ The researchers suggest that glycosaminoglycans, whose binding to TREM2 is reduced by the risk variant R47H, might play a role in the difference in cellular signaling. The same study also suggested that the basic and hydrophobic patches on the extracellular domains in their crystal structure could mediate binding with phospholipids.^31^ Further structural studies will need to be done to determine the molecular mechanisms of TREM2-ligand interactions and how they affect the activation of signaling pathways.

### TREM2 Signaling

The most important transmembrane adapter for TREMs is the DNAX-activating protein of 12 kDa (DAP12).^5^ DAP12 contains an intracellular immunoreceptor tyrosine-based activation motif (ITAM). Upon TREM-DAP12 activation, SRC kinases phosphorylate tyrosines on the ITAMs, allowing for the ligation of spleen-associated tyrosine kinase (SYK) to the phosphorylated ITAM.^5, 32^ The SYK activates downstream signaling molecules such as phosphatidylinositol 3-kinase (PI3K) and phospholipase C gamma 2 (PLCγ2).^5^ PI3K activates the AKT kinase, which triggers the mammalian target of rapamycin (mTOR) pathway to initiate.^32^ The mTOR pathway promotes cell survival and proliferation by supporting ATP production and protein synthesis.^33, 34^ Specifically, the mTOR complex 1 (mTORC1) phosphorylates the eukaryotic translation initiation factor 4E (eIF4E)-binding protein 1 (4E-BP1).^33, 35^ This inhibits 4E-BP1, which in its hypophosphorylated (uninhibited) form, binds to eIF4E, preventing eIF4E from binding to eIF4G and forming the eIF4F complex, which is responsible for recruiting 40S ribosomal subunit to the 5′-cap of mRNAs.^35, 36^ This means that mTORC1 prevents 4E-BP1 from inhibiting translation. mTORC1 has been shown to selectively promote the translation of nucleus-encoded mitochondria-related mRNAs, such as those for the subunits of the oxidative phosphorylation Complexes I and V, which can lead to an increase in mitochondrial ability to produce ATP, the energy from which can facilitate further protein synthesis necessary for cell survival and proliferation.^33^ Detailed signaling pathway for TREM2 is outlined below. Refer to Fig. 1.

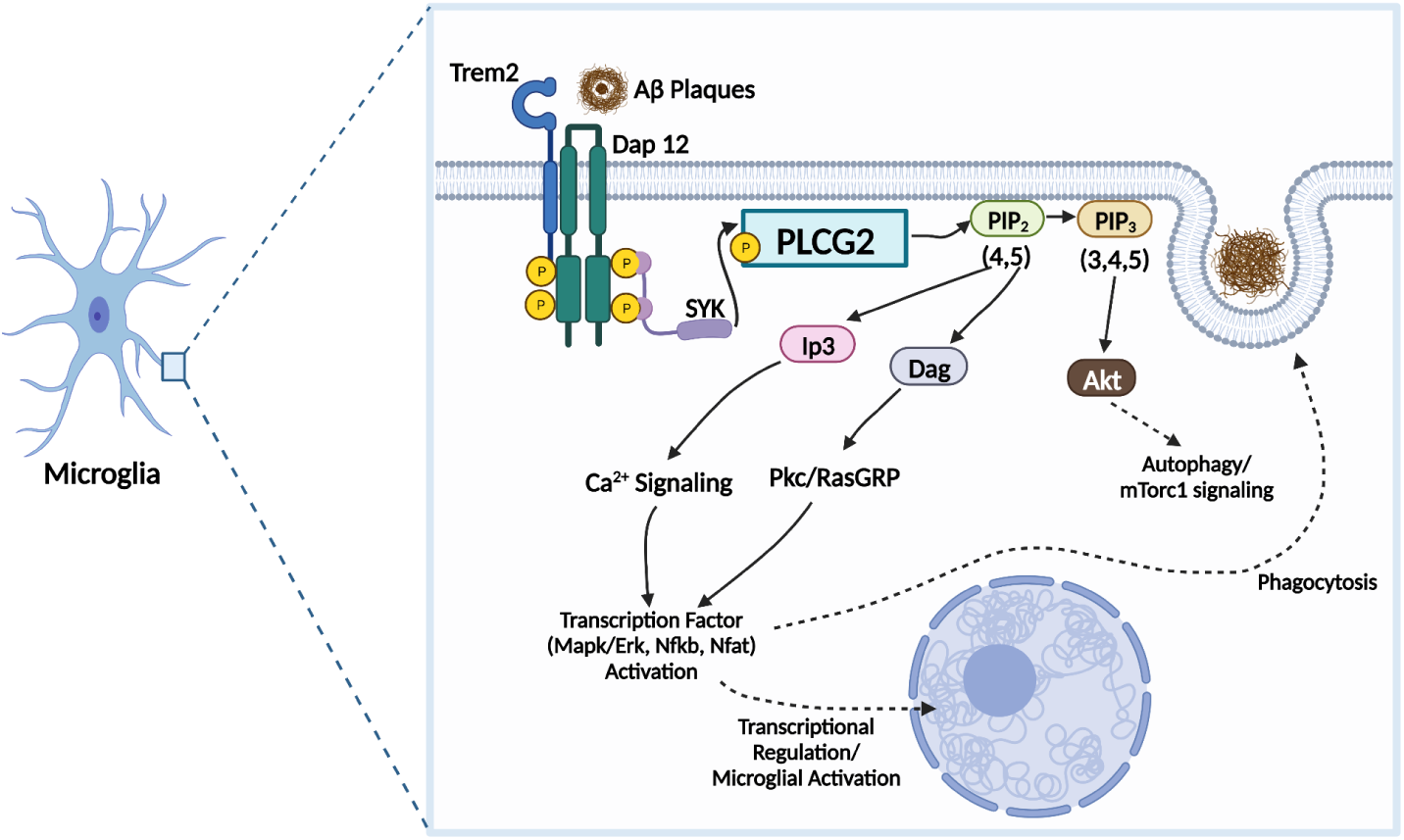

In a study on TREM2^−/−^ 5XFAD mice and post-mortem brain samples from AD patients heterozygous for R47H and R62H, it was found that they contained microglia abundant in vesicular structures and which were LC3+.^34^ LC3+ is a commonly used indication of autophagosomes, which are vesicles participating in cellular autophagy, the process by which macromolecules, pathogens, and organelles are degraded.^37^ While autophagy can occur to eliminate harmful misfolded proteins, aggregate-prone proteins, and damaged organelles, it can also occur due to cellular stress from cells having insufficient energy or inputs for protein synthesis.^34, 37, 38^ This means that R47H likely leads to reduced mTOR signaling, diminishing ATP production and protein synthesis leading to fewer microglia surviving and proliferating in response to Aβ accumulation.

In addition to the PI3K-AKT-mTOR pathway, SYK also activates PLCγ2, which cleaves phosphatidylinositol 4,5-bisphosphate (PIP2) into inositol 1,4,5-trisphosphate (IP3) and diacylglycerol (DAG).^5, 39^ IP3 triggers ligand-gated ion channels on the endoplasmic reticulum to release Ca^2+^.^39^ Calcium, as a common secondary messenger, can go on to trigger many other signaling pathways. For instance, increased intracellular calcium triggers calcineurin to dephosphorylate the nuclear factor of activated T cells (NFAT), allowing it to enter the nucleus.^40, 41^ NFAT is a transcription factor that plays a role in the transcription of various genes in many cell types.^42^ In immune cells such as microglia, it helps facilitate the expression of cytokines. One study using in vitro microglia and an AD mouse model determined, through the use of NFAT inhibitors FK506 and tat-VIVIT, that NFAT inhibition decreased secretion of cytokines TNF-α and interleukin-6 in vitro following stimulation with Aβ or LPS.^41^ However, the researchers also found that those same cytokine concentrations were significantly diminished by the NFAT inhibitors in the mouse models’ spleens but not brains (they attributed this to insufficient dosage). This study also found reduced Aβ plaque burden in the mouse models when treated with the NFAT inhibitors and that in vitro microglia had increased phagocytic uptake of fibrillar Aβ while treated with FK506.^41^ These findings imply that NFAT’s pro-inflammatory cytokine expression can be suppressed without diminishing microglial ability to eliminate Aβ.

Increased intracellular calcium due to IP3 activity, in addition to mediating further signaling pathways, also plays an essential function in phagocytosis by triggering the activity of contractile proteins in the cytoskeleton, such as myosin and actin.^43^ DAG produced from the cleavage of PIP2 triggers protein kinase C (PKC) and Ras guanyl-nucleotide-releasing proteins (RasGRPs), which subsequently activate Nuclear Factor-kappa B (NF-κB) and mitogen-activated protein kinase (MAPK) pathways.^39^ NF-κB promotes the expression of multiple pro-inflammatory factors, including cytokines, chemokines, cell cycle regulators, anti-apoptotic factors, and adhesion molecules.^44^ This allows microglia and other immune cells to survive, proliferate, enhance the inflammatory response, and migrate into tissues to eliminate pathogens or other harmful substances.^44, 45^

The MAPK pathway is a crucial signaling pathway in cell growth, development, differentiation, proliferation, and apoptosis.^46^ The dysregulation of MAPK signaling is heavily implicated in supporting tumorigenesis. MAPK signaling cascades generally share a motif of three protein kinases called MAPK kinase kinase (MAP3K), MAPK kinase (MAPKK), and MAPK, in addition to variable upstream and downstream signaling molecules.^46^ The best-understood pathway is the MAPK/Extracellular signal-regulated kinase (ERK) pathway, which consists of Ras (upstream activator), Raf (MAP3K), MEK1/2 (MAPKK), and ERK1/2 (MAPK). Ras binds to Raf-1, causing it to translocate from the cytoplasm to the cell membrane. Raf then phosphorylates MEK, which then phosphorylates ERK. Phosphorylated ERK can then trigger other protein kinases, phosphorylate target proteins in the cytoplasm, or dimerize and enter the nucleus to phosphorylate and, thus, activate various transcription factors.^46^ In the mechanism outlined in this paper, RasGRPs, stimulated by DAG, are responsible for converting Ras GDP (the inactive form) to Ras GTP (the active form).^47^ However, it is important to note that other mechanisms, including those using other Ras guanine nucleotide exchange factors (Ras GEFs), such as son of sevenless (SOS) proteins, are also likely to be involved in TREM2-ERK signaling.

#### Neurofilament Light Chain

Neurofilament is a cytoskeletal protein found only in neurons, specifically their axons. It has four subunits: light, medium, heavy, and alpha-internexin, with NFL being the most abundant. NFL is a biomarker in other diseases, such as frontotemporal and vascular dementia, with increased CSF NFL levels indicative of neurodegeneration.^48^ It has also been shown to be a potentially useful biomarker for AD. In a study by Giacomucci et al., plasma NFL levels were compared with CSF biomarkers and neuropsychological scores to identify if NFL correlated with those indicators and could help diagnose AD in its early stages.^49^ Patients involved include 23 with AD, 53 with mild cognitive impairment (MCI), and 34 with subjective cognitive decline (SCD). They were scored using the mini-mental state examination and other neuropsychological measurements and received plasma NFL analysis, CSF tau and phosphorylated tau (p-tau) measurements, and CSF or PET analysis of amyloid burden.^49^ The results were that amyloid-positive MCI patients had NFL levels similar to AD but higher than MCI amyloid-negatives, who in turn had NFL levels comparable to those of SCD amyloid-positives. NFL correlated directly with p-tau and total tau (t-tau) in the MCI group, inversely with CSF Aβ42 and Aβ42/Aβ40 ratio in the MCI group, directly with phosphorylated tau in the SCD group, and inversely with memory test scores.^49^ Additionally, there are “no correlations between NfL levels and CSF biomarkers in AD patients,” leading the researchers to suggest that neurodegeneration plateaus at some point, thus no longer being correlated with the other biomarkers. In summary, this study indicates that NFL can serve as a valuable biomarker for preclinical and prodromal AD but can become less reliable afterward.

A meta-analysis by Jin et al. finds similar results, such as a positive correlation between CSF NFL and CSF t-tau, p-tau, neurogranin, and YLK-40. However, unlike Giacomucci et al., this analysis finds that CSF NFL and Aβ are not correlated, rather than inversely correlated.^48, 49^ They also note that “The presence of more amyloid plaques leads to lower Aβ levels in CSF, whereas higher neurofibrillary tangles cause higher tau CSF levels.” Considering the combined results of these papers, neurodegeneration might not plateau after its initial inverse progression with Aβ, but rather, the correlation between its marker (NFL) and Aβ ends at some point, such as when Aβ deposition plateaus, as demonstrated by Engler et al.^12^ In conclusion, NFL correlates positively with some CSF biomarkers, but its relationship with Aβ levels is less clear. Nevertheless, it is likely a useful clinical marker for diagnosing AD and predicting its prognosis.

## Methods

In order to better understand the mechanism for which TREM2 alteration affects the genetic and phenotypic changes in the 5xFAD mice model, MODEL-AD Database by Sage Bionetworks was used. The MODEL-AD database contained the biomarker concentration in the cerebral cortex and hippocampus of 4 month and 12 month 5xFAD TREM2^R47H^ mice. A portion of the dataset is below. Refer to Fig. 2.

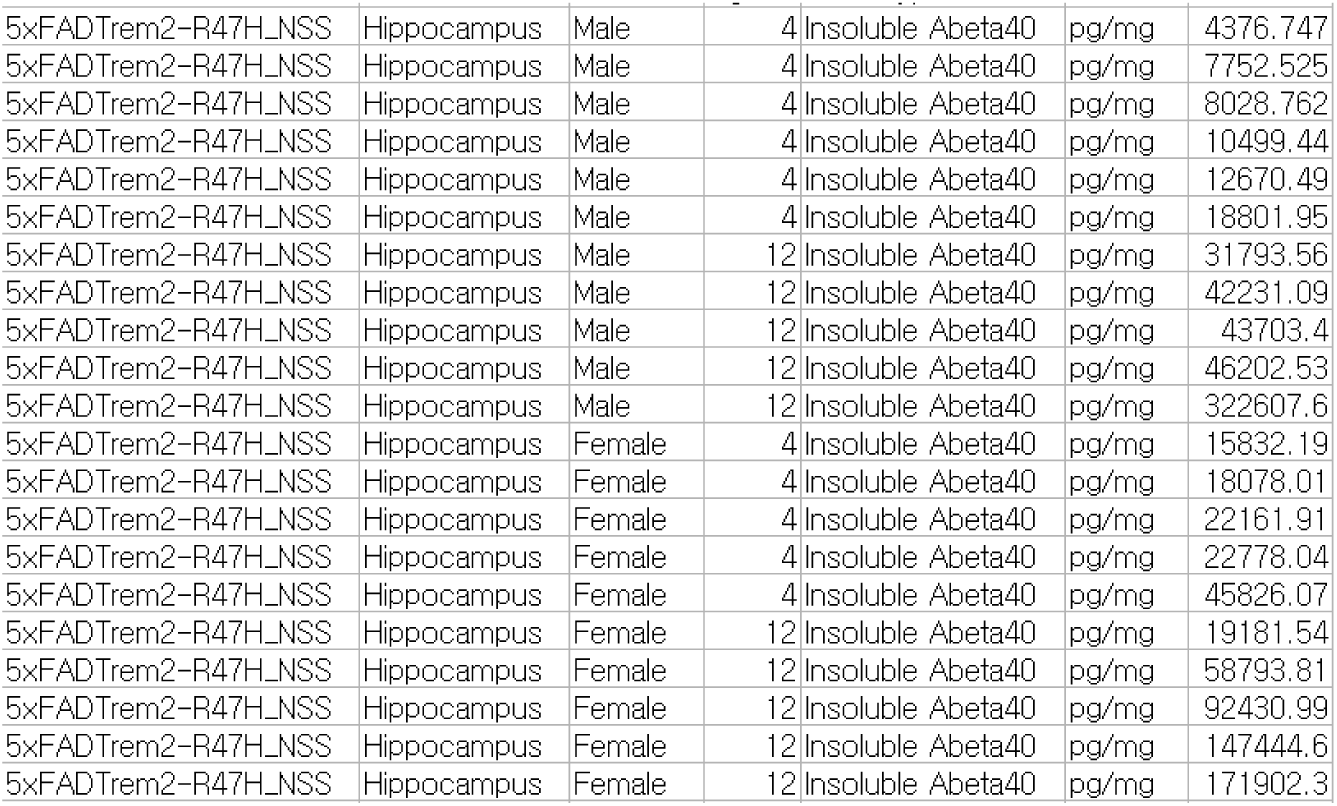

Furthermore, the MODEL-AD database also contained the regulatory status of various genes within the brains of 4,8,12, and 24 month TREM2^-/-^ 5xFAD mice. A portion of that dataset is shown below. The genes that were not yet studied were highlighted in yellow. Refer to Fig. 3.

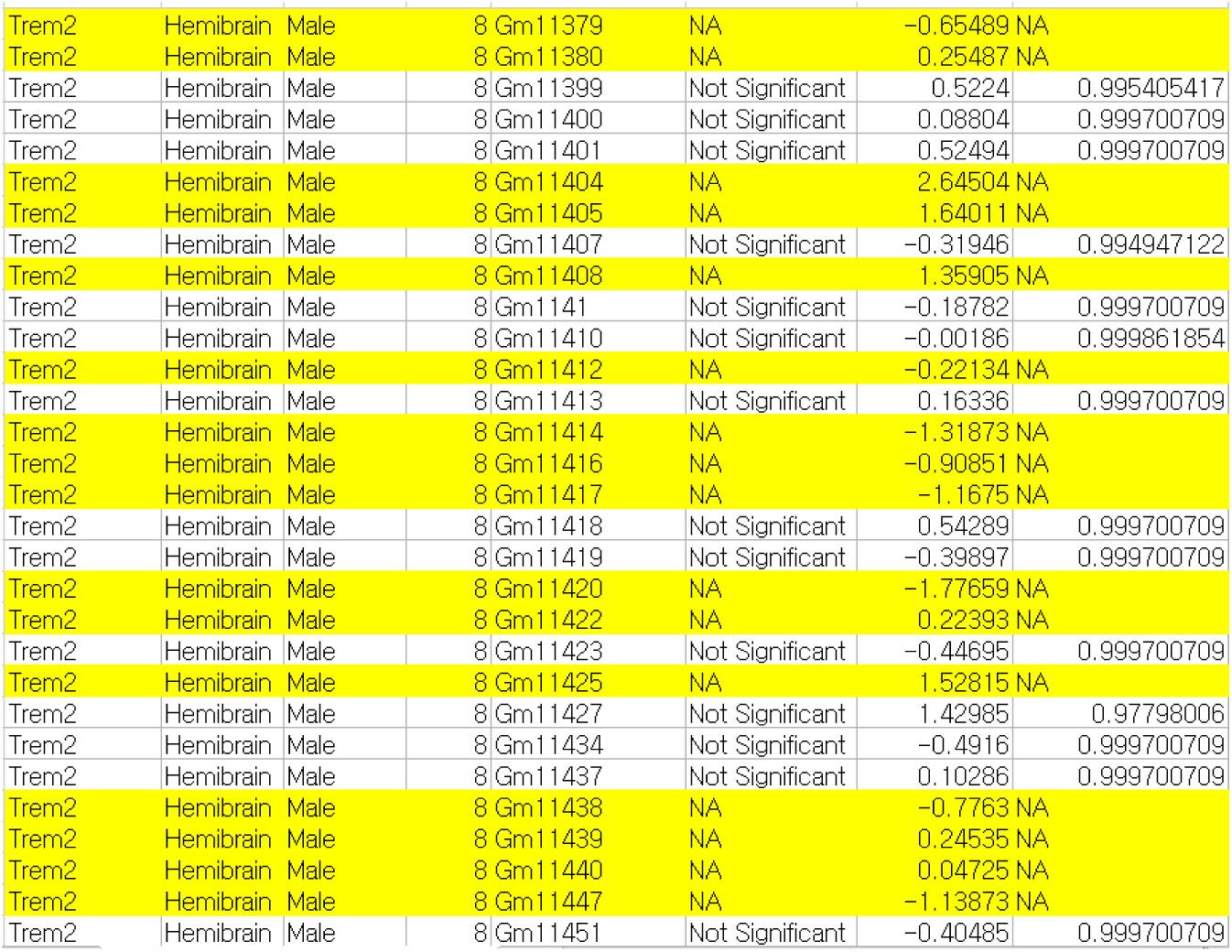

The average of biomarker concentration were calculated and analyzed for percentage increase of biomarkers as the mice aged from 4 month and 12 month. The gene regulatory status dataset were analyzed for any genes that were upregulated and downregulated. Then these results were normalized based on the proportion of the genes that were not studied. Detailed methodology outline is shown below. Finally, images of TREM2^-/-^ and WT of 5xFAD mice cortex tissue stained with X34 and LAMP1 were taken with Leica Confocal Microscopy. X34 is a stain that marks the β-pleated sheets within Aβ plaques. Lysosomal associated membrane protein1 (LAMP1) is a stain that marks for neuronal dystrophy as lysosomes serve as degradation hubs for autophagic components.^50^ Refer to Fig. 4.

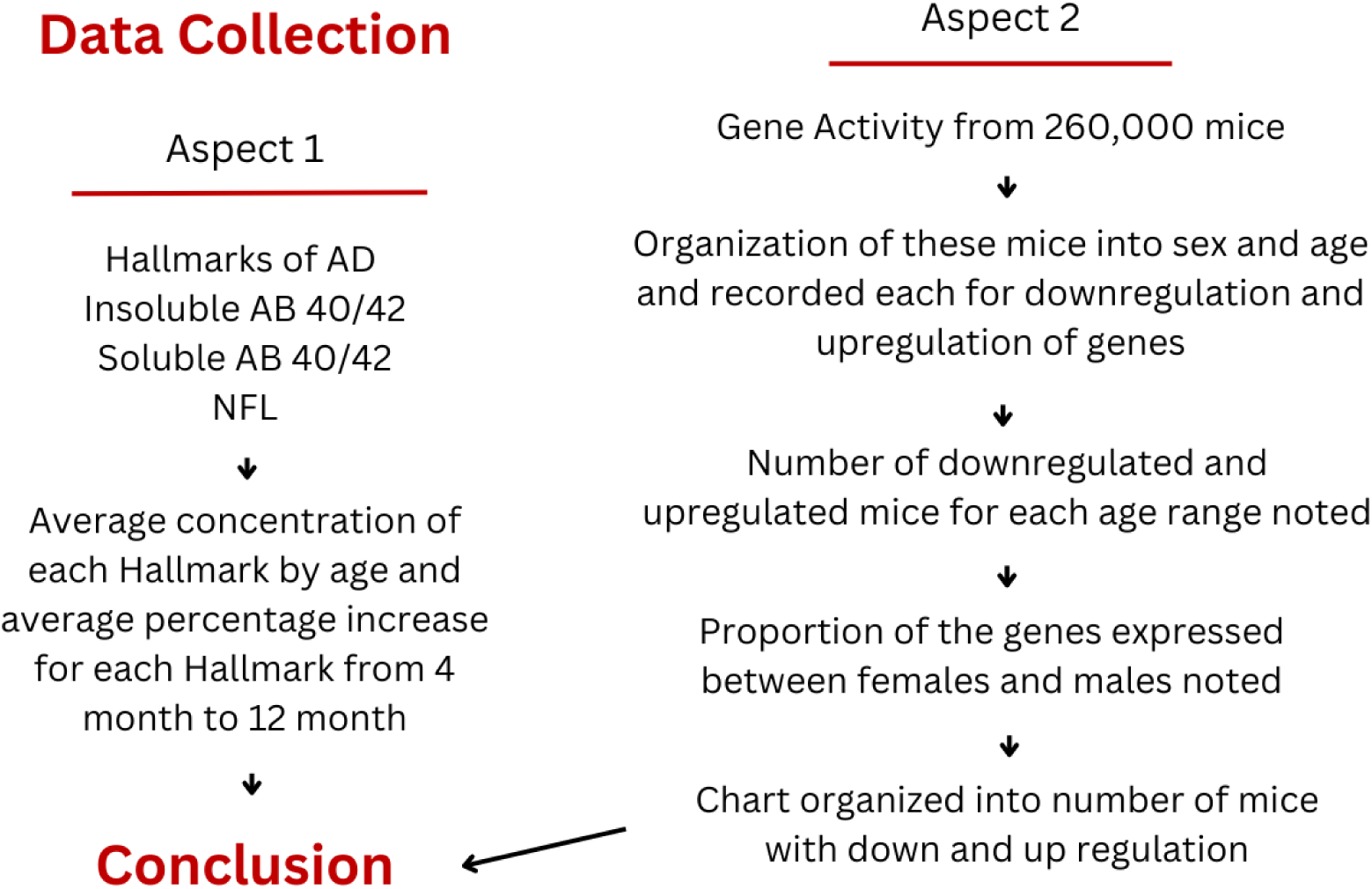

## Results

Below were the results of the average biomarker concentration of 4 month and 12 month 5xFAD TREM2^R47H^ mice model. These datasets were further analyzed for percentage increase of biomarkers as the mice aged from 4 months and 12 months. Refer to Fig. 5. Refer to Fig. 6.

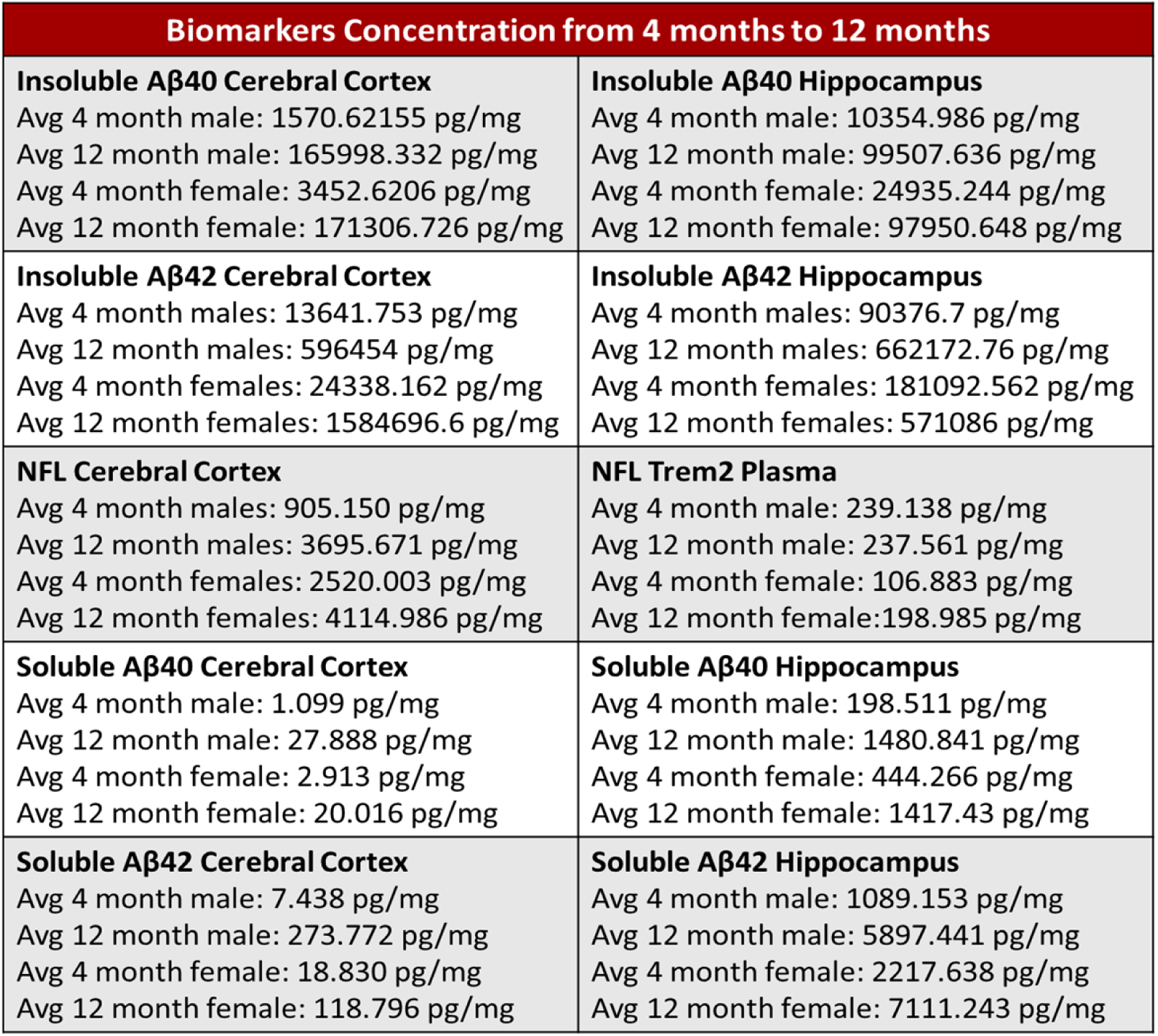

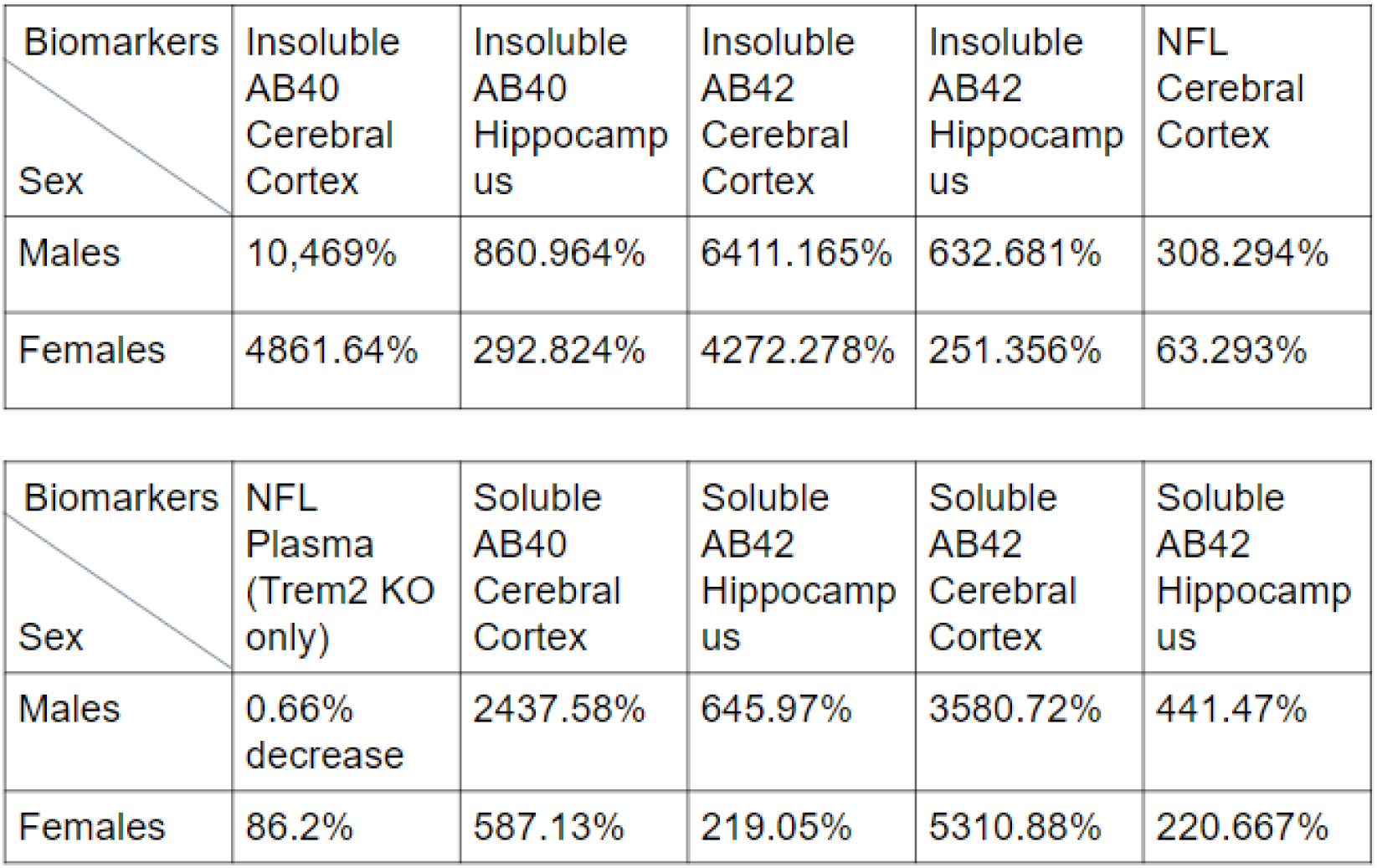

The females generally had lower percentage increases, which indicates that for the 5xFAD TREM2^-/-^ mice models, the females had larger values of AD biomarkers as compared to the males before both sexes began to display similar concentration of biomarkers at around 12 months and onwards.

However, a major discrepancy came from NFL plasma data where females had a larger percentage increase. NFL plasma data were derived from TREM2^-/-^ mice, which shows a clear sign that 5xFAD increases concentration of biomarkers in females more than males. This might show a differential mechanism between the TREM2 alteration affecting the biomarker concentration sex differences in 5xFAD mice.

The results for gene regulatory status analysis were shown below. The proportion of genes analyzed were multiplied by the total number of genes on the database by each age and sex. Downregulation occurred in 12 month mice and 24 month mice. In 12 months aged mice, the ENSMUSG family of gene, RP 23 family of genes, RP 24 family of genes were downregulated. In 24 months aged mice, Gimap3, Gpnmb, Cxcl16, Ccl family of genes, and Cd family of genes were downregulated. Upregulation occurred in 4,12,and 24 month mice. In 4 months aged mice, Gm28196 gene was upregulated and continued to be upregulated. In 12 month aged mice, Gm12756, Gm3239 genes, and the Hoxa, Hoxb, Hoxc family of genes were upregulated. In 24 month aged mice, Gm family of genes and the ENMUSG family of genes were upregulated. Refer to Fig. 7.

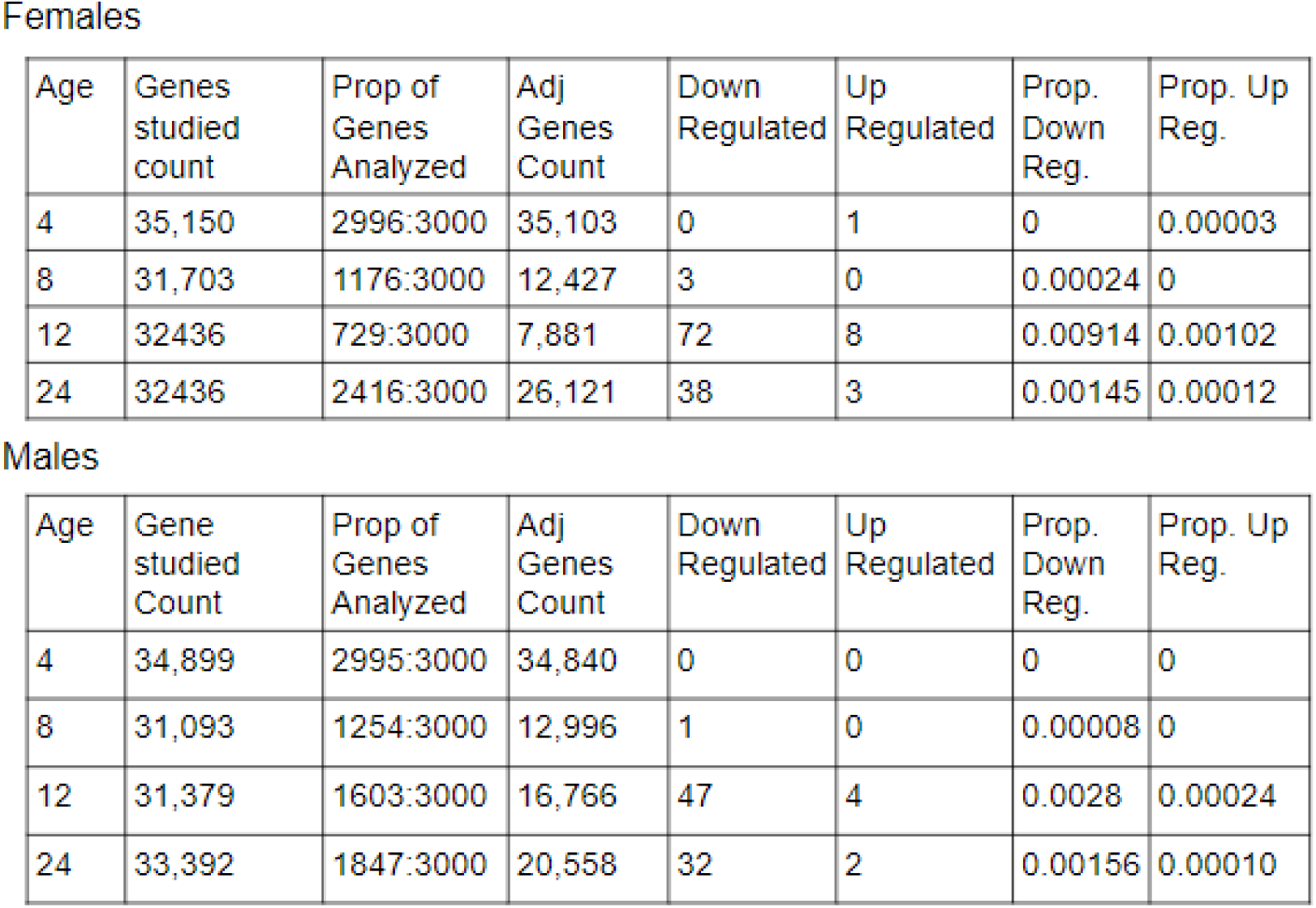

Finally, the confocal images of WT and TREM2^-/-^ 5xFAD mice were shown below. X34 stains for plaques were marked with blue while LAMP1 stains for neuron damage were marked with green. Confocal images allows each layer of multistack (z-stack) images to be of the same focus and rejects any out of focus images from each layer of z-stack which helps it produce high resolution images.^51^

The qualitative analysis of these images show that there is significantly more presence of neuron damage and death with the TREM2^-/-^ mice models, highlighting TREM2 genes function of neuronal protection. Refer to Fig. 8.

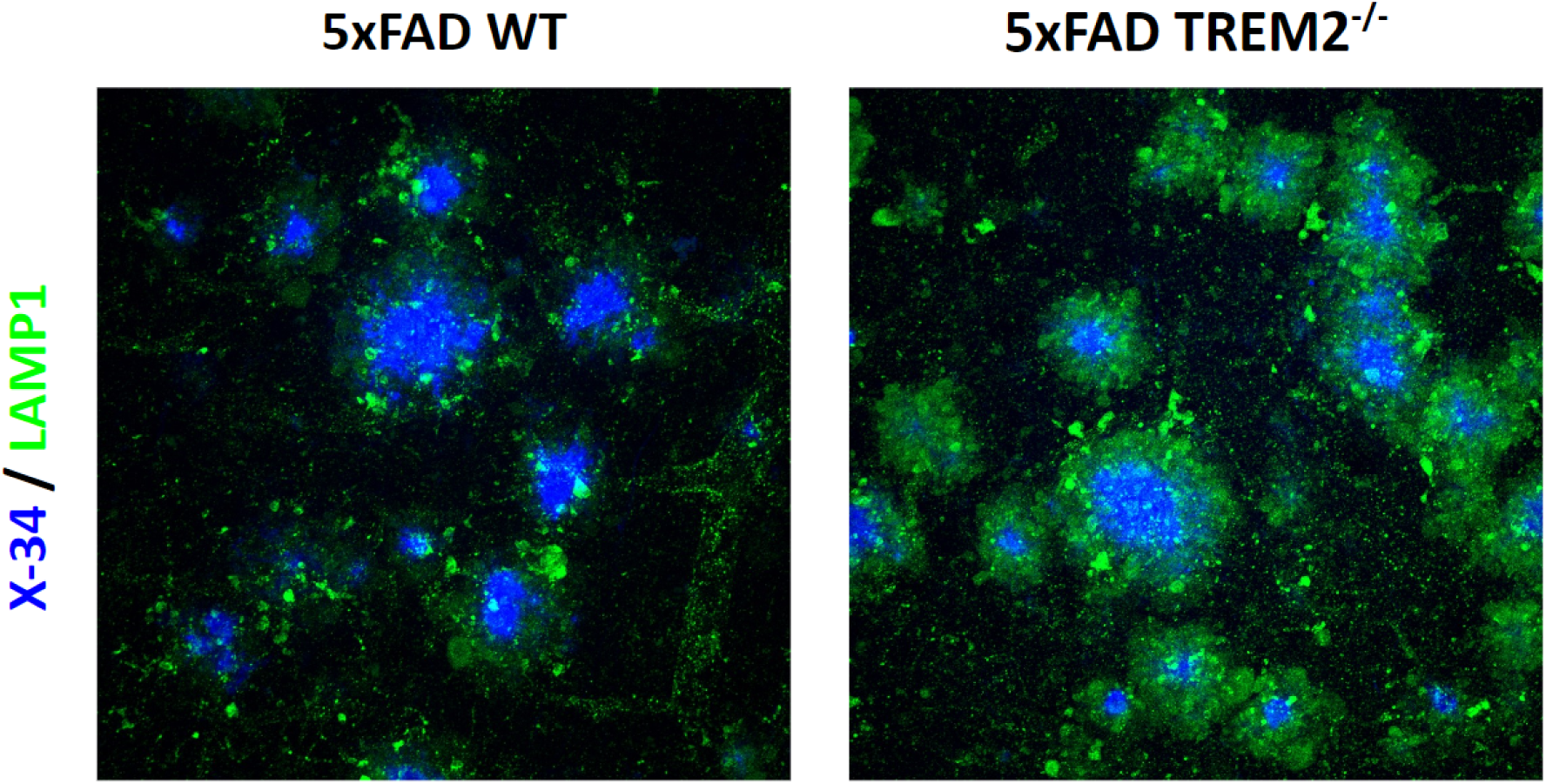

Then, the percentage area of both X34 and LAMP1 stains were analyzed on ImageJ and the data points for each cross section of the image stack were further organized into PRISMS software. The results for that are below. Refer to Fig. 9.

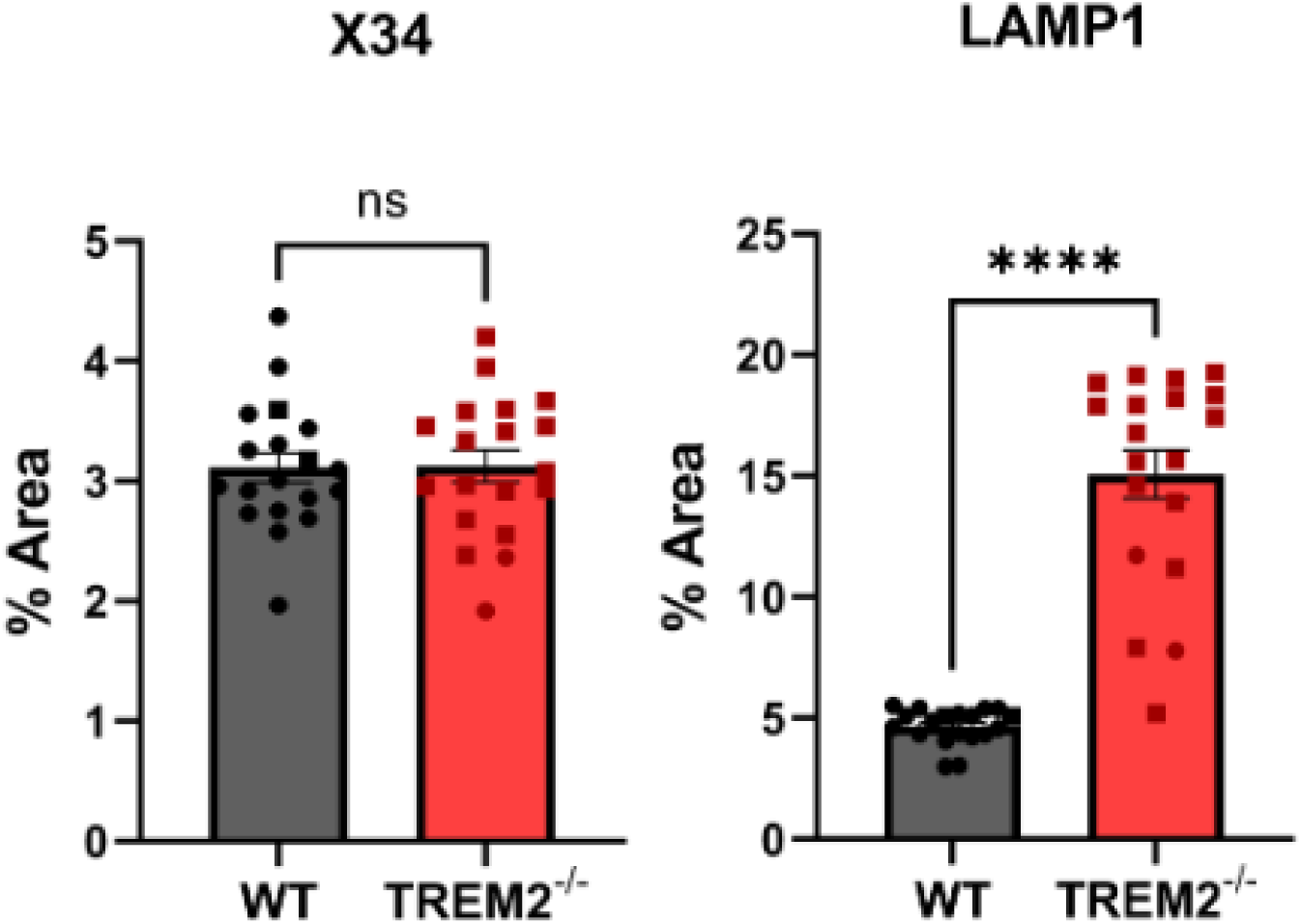

The plaque percentage area differences between the two genotypes were minimal, however, there was very significant evidence of increased neuronal damage with TREM2^-/-^. This shows that the plaque concentration did not have any sort of bearing on the neuronal changes that occurred and isolates the cause to the genotypic difference. These results confirm that TREM2 helps protect neurons from plaque toxicity by quickly activating the mechanisms within the brain to counteract the plaques.

## Limitation

As a significant proportion of the genes from each age and sex of the TREM2^-/-^ were not studied, this shows that with the update of the MODEL-AD database and more genes being studied, there might be significant deviation from the current results.

## Conclusions

TREM2^R47H^ mutated 5xFAD mice have been shown to have initial sex differences in biomarker concentration that equalized in the later ages (12 months). However, it is not known if this characteristic of biomarker difference is caused by the TREM2^R47H^ mutation or if this is the characteristic of 5xFAD mice models.

TREM2^-/-^ 5xFAD mice are shown to be associated with several genotypic regulation and therefore means that TREM2 gene is associated with these genes. Confocal image analysis of 7.5 month 5xFAD TREM2^-/-^ mice show that the TREM2 receptor presence significantly lowers the neuronal damage rate.

Overall. These results further the knowledge of TREM2 gene mechanisms through MODEL-AD analysis and Confocal Imaging methods.

## References

[1] Gauthier S, Webster C, Servaes S, et al. World Alzheimer Report 2022: Life after diagnosis: Navigating treatment, care and support. London, England: Alzheimer’s Disease International. 2022. Accessed July 20,2023

[2] World Health Organization. The top 10 causes of death. WHO | World Health Organization. December 9, 2020. Accessed July 20, 2023 https://www.who.int/news-room/fact-sheets/detail/the-top-10-causes-of-death

[3] Guerreiro R, Wojtas A, Bras J, et al. TREM2 Variants in Alzheimer’s Disease . The New England Journal of Medicine. January 10, 2013. Accessed July 24, 2023. https://www.nejm.org/doi/10.1056/NEJMoa1211851.

[4] Jonsson T, Stefansson H, Steinberg S, Jonsdottir I, Jonsson P. Variant of TREM2 Associated with the Risk of Alzheimer’s Disease. The New England Journal of Medicine. January 10, 2013. Accessed August 5, 2023. https://www.nejm.org/doi/10.1056/NEJMoa1211103.

[5] Colonna M. The biology of Trem Receptors. Nature News. February 7, 2023. Accessed August 5, 2023. https://www.nature.com/articles/s41577-023-00837-1.

[6] Chen G, Xu T, Yan Y, et al. Amyloid beta: Structure, biology and structure-based therapeutic development. Nature News. July 17, 2017. Accessed August 5, 2023. https://www.nature.com/articles/aps201728.

[7] Kametani F, Hasegawa M. Reconsideration of Amyloid Hypothesis and Tau Hypothesis in Alzheimer’s Disease. Frontiers in neuroscience. January 30, 2018. Accessed August 5, 2023. https://www.ncbi.nlm.nih.gov/pmc/articles/PMC5797629/.

[8] Jeong H, Shin H, Hong S, Kim Y. Physiological roles of monomeric amyloid-β and implications for alzheimer’s disease therapeutics. Experimental neurobiology. April 30, 2022. Accessed August 5, 2023. https://www.ncbi.nlm.nih.gov/pmc/articles/PMC9194638/.

[9] Li X, Yang T, Sun Z. Hormesis in health and chronic diseases. Trends in endocrinology and metabolism: TEM. December 2019. Accessed August 5, 2023. https://www.ncbi.nlm.nih.gov/pmc/articles/PMC6875627/.

[10] Priller C, Bauer T, Mitteregger G, Krebs B, Kretzschmar HA, Herms J. Synapse formation and function is modulated by the amyloid precursor protein. The Journal of neuroscience : the official journal of the Society for Neuroscience. July 5, 2006. Accessed August 5, 2023. https://www.ncbi.nlm.nih.gov/pmc/articles/PMC6673945/.

[11] Wang J, Dickson DW, Trojanowski JQ, Lee V. The Levels of Soluble versus Insoluble Brain Aβ Distinguish Alzheimer’s Disease from Normal and Pathologic Aging. ScienceDirect. 1999. Accessed August 5, 2023. 10.1006/exnr.1999.7085.

[12] Engler H, Forsberg A, Almkvist O, et al. Two-year follow-up of amyloid deposition in patients with Alzheimer’s disease. OUP Academic. July 19, 2006. Accessed August 5, 2023. https://academic.oup.com/brain/article/129/11/2856/289548.

[13] Yamin G, Teplow DB. Pittsburgh compound-B (PIB) binds amyloid β-protein protofibrils. Journal of neurochemistry. January 2017. Accessed August 5, 2023. https://www.ncbi.nlm.nih.gov/pmc/articles/PMC5225051/.

[14] Maarouf CL, Daugs ID, Kokjohn TA, et al. Alzheimer’s disease and non-demented high pathology control nonagenarians: Comparing and contrasting the biochemistry of cognitively successful aging. PloS one. 2011. Accessed August 5, 2023. https://www.ncbi.nlm.nih.gov/pmc/articles/PMC3210154/.

[15] Morris GP, Clark IA, Vissel B. Inconsistencies and controversies surrounding the amyloid hypothesis of alzheimer’s disease. Acta neuropathologica communications. September 18, 2014. Accessed August 5, 2023. https://www.ncbi.nlm.nih.gov/pmc/articles/PMC4207354/.

[16] Howard Jay Aizenstein M. Frequent amyloid deposition without significant cognitive impairment among the elderly. Archives of Neurology. November 10, 2008. Accessed August 5, 2023. https://jamanetwork.com/journals/jamaneurology/fullarticle/1107509.

[17] Murphy MP, LeVine H. Alzheimer’s Disease and the β-Amyloid Peptide. Journal of Alzheimer’s disease : JAD. 2010. Accessed August 5, 2023. https://www.ncbi.nlm.nih.gov/pmc/articles/PMC2813509/.

[18] Holmes C, Boche D, Wilkinson D, Yadegarfar G. Long-term effects of Aβ42 immunisation in Alzheimer’s disease: follow-up of a randomised, placebo-controlled phase I trial. The Lancet. July 19, 2008. Accessed August 5, 2023. https://www.thelancet.com/journals/lancet/article/PIIS0140-6736(08)61075-2/fulltext.

[19] JA N, M L, C H, et al. AN-1792. ALZFORUM. 2004. Accessed August 5, 2023. https://www.alzforum.org/therapeutics/an-1792.

[20] Goure WF, Krafft GA, Jerecic J, Hefti F. Targeting the proper amyloid-beta neuronal toxins: a path forward for Alzheimer’s disease immunotherapeutics. BioMed Central. July 9, 2014. Accessed August 5, 2023. https://alzres.biomedcentral.com/articles/10.1186/alzrt272.

[21] Shankar GM, Li S, Mehta TH, et al. Amyloid-beta protein dimers isolated directly from Alzheimer’s brains impair synaptic plasticity and memory. Nature medicine. August 2008. Accessed August 5, 2023. https://www.ncbi.nlm.nih.gov/pmc/articles/PMC2772133/.

[22] Nabavi S, Fox R, Proulx CD, Lin JY, Tsien RY, Malinow R. Engineering A memory with ltd and LTP. Nature. July 17, 2014. Accessed August 5, 2023. https://www.ncbi.nlm.nih.gov/pmc/articles/PMC4210354/.

[23] Ge Y, Dong Z, Bagot RC, et al. Hippocampal long-term depression is required for the consolidation of Spatial Memory. Proceedings of the National Academy of Sciences of the United States of America. September 21, 2010. Accessed August 5, 2023. https://www.ncbi.nlm.nih.gov/pmc/articles/PMC2944752/.

[24] Kim J, Chakrabarty P, Hanna A, et al. Normal cognition in transgenic BRI2-Aβ mice. Molecular neurodegeneration. May 12, 2013. Accessed August 5, 2023. https://www.ncbi.nlm.nih.gov/pmc/articles/PMC3658944/.

[25] Takahashi K, Rochford CDP, Neumann H. Clearance of apoptotic neurons without inflammation by microglial triggering receptor expressed on myeloid cells-2. Journal of Experimental Medicine. February 21, 2005. Accessed August 5, 2023. https://rupress.org/jem/article/201/4/647/52887/Clearance-of-apoptotic-neurons-without.

[26] Popko K, Gorska E, Stelmaszczyk-Emmel A, et al. Proinflammatory cytokines IL-6 and TNF-α and the development of inflammation in obese subjects. European journal of medical research. November 4, 2010. Accessed August 5, 2023. https://www.ncbi.nlm.nih.gov/pmc/articles/PMC4360270/.

[27] Zamora R, Vodovotz Y, Billiar TR. Inducible nitric oxide synthase and inflammatory diseases - molecular medicine. BioMed Central. May 1, 2000. Accessed August 5, 2023. https://molmed.biomedcentral.com/articles/10.1007/BF03401781.

[28] Picón-Pagès P, Garcia-Buendia J, Muñoz F. Functions and dysfunctions of nitric oxide in brain. Biochimica et Biophysica Acta (BBA) - Molecular Basis of Disease. November 27, 2018. Accessed August 5, 2023. https://www.sciencedirect.com/science/article/pii/S0925443918304526.

[29] Takahashi T, Suzuki T. Role of sulfatide in normal and pathological cells and tissues. Journal of lipid research. August 2012. Accessed August 5, 2023. https://www.ncbi.nlm.nih.gov/pmc/articles/PMC3540844/.

[30] Zhao Y, Wu X, Li X, An Z. TREM2 Is a Receptor for β-Amyloid that Mediates Microglial Function. Neuron. January 31, 2018. Accessed August 5, 2023. https://www.cell.com/neuron/fulltext/S0896-6273(18)30056-4.

[31] Kober DL, Alexander-Brett JM, Karch CM, et al. Neurodegenerative disease mutations in TREM2 reveal a functional surface and distinct loss-of-function mechanisms. eLife. December 20, 2016. Accessed August 5, 2023. https://elifesciences.org/articles/20391.

[32] Hall-Roberts H, Agarwal D, Obst J, et al. TREM2 Alzheimer’s variant R47H causes similar transcriptional dysregulation to knockout, yet only subtle functional phenotypes in human iPSC-derived macrophages. BioMed Central. November 16, 2020. Accessed August 5, 2023. https://alzres.biomedcentral.com/articles/10.1186/s13195-020-00709-z.

[33] Morita M, Gravel S-P, Chénard V, Sikström K, Zheng L, Alain T. MTORC1 controls mitochondrial activity and biogenesis through 4E-bp-dependent translational regulation. Cell Metabolism. November 5, 2013. Accessed August 5, 2023. https://www.sciencedirect.com/science/article/pii/S1550413113004130.

[34] Ulland T, Song W, Huang S, Artyomov M. TREM2 Maintains Microglial Metabolic Fitness in Alzheimer’s Disease. Cell. July 23, 2017. Accessed August 5, 2023. https://www.cell.com/cell/fulltext/S0092-8674(17)30830-9.

[35] Qin X, Jiang B, Zhang Y. 4E-BP1, a multifactor regulated multifunctional protein. Cell cycle (Georgetown, Tex.). 2016. Accessed August 5, 2023. https://www.ncbi.nlm.nih.gov/pmc/articles/PMC4845917/.

[36] UniProt. Q62622 · 4EBP1_RAT. UniProt. 2018. Accessed August 5, 2023. https://www.uniprot.org/uniprotkb/Q62622/entry.

[37] Runwal G, Stamatakou E, Siddiqi FH, Puri C, Zhu Y, Rubinsztein DC. LC3-positive structures are prominent in autophagy-deficient cells. Nature News. July 12, 2019. Accessed August 5, 2023. https://www.nature.com/articles/s41598-019-46657-z.

[38] Rabinowitz JD, White E. Autophagy and metabolism. Science (New York, N.Y.). December 3, 2010. Accessed August 5, 2023. https://www.ncbi.nlm.nih.gov/pmc/articles/PMC3010857/.

[39] Obst J, Hall-Roberts HL, Smith TB, et al. PLCγ2 regulates TREM2 signaling and integrin-mediated adhesion and migration of human iPSC-derived macrophages. Scientific reports. October 6, 2021. Accessed August 5, 2023. https://www.ncbi.nlm.nih.gov/pmc/articles/PMC8494732/.

[40] Rojanathammanee L, Floden AM, Manocha GD, Combs CK. Attenuation of microglial activation in a mouse model of Alzheimer’s disease via NFAT inhibition. BioMed Central. March 4, 2015. Accessed August 5, 2023. https://jneuroinflammation.biomedcentral.com/articles/10.1186/s12974-015-0255-2.

[41] Creamer TP. Calcineurin - cell communication and signaling. BioMed Central. August 28, 2020. Accessed August 5, 2023. https://biosignaling.biomedcentral.com/articles/10.1186/s12964-020-00636-4.

[42] Horsley V, Pavlath GK. NFAT: Ubiquitous regulator of cell differentiation and adaptation. The Journal of cell biology. March 4, 2002. Accessed August 5, 2023. https://www.ncbi.nlm.nih.gov/pmc/articles/PMC2173310/.

[43] Melendez A. Calcium Signaling During Phagocytosis. National Library of Medicine. 2013. Accessed August 5, 2023. https://www.ncbi.nlm.nih.gov/books/NBK5971/.

[44] Liu T, Zhang L, Joo D, Sun S-C. NF-ΚB signaling in inflammation. Nature News. July 14, 2017. Accessed August 5, 2023. https://www.nature.com/articles/sigtrans201723.

[45] Ren G, Roberts AI, Shi Y. Adhesion molecules: Key players in mesenchymal stem cell-mediated immunosuppression. Cell adhesion & migration. 2011. Accessed August 5, 2023. https://www.ncbi.nlm.nih.gov/pmc/articles/PMC3038091/.

[46] Guo Y-J, Pan W-W, Liu S-B, Shen Z-F, Xu Y, Hu L-L. Erk/MAPK signalling pathway and tumorigenesis. Experimental and therapeutic medicine. March 2020. Accessed August 5, 2023. https://www.ncbi.nlm.nih.gov/pmc/articles/PMC7027163/.

[47] Jun JE, Rubio I, Roose JP. Regulation of Ras exchange factors and cellular localization of Ras activation by lipid messengers in T cells. Frontiers. August 2, 2013. Accessed August 5, 2023. https://www.frontiersin.org/articles/10.3389/fimmu.2013.00239/full.

[48] Jin M, Cao L, Dai Y. Role of Neurofilament Light Chain as a Potential Biomarker for Alzheimer’s Disease: A Correlative Meta-Analysis. Frontiers. August 27, 2019. Accessed August 5, 2023. https://www.frontiersin.org/articles/10.3389/fnagi.2019.00254/full.

[49] Giacomucci G, Mazzeo S, Bagnoli S, et al. Plasma neurofilament light chain as a biomarker of Alzheimer’s disease in subjective cognitive decline and mild cognitive impairment. Journal of neurology. August 2022. Accessed August 5, 2023. https://www.ncbi.nlm.nih.gov/pmc/articles/PMC9293849/.

[50] Cheng X-T, Xie Y-X, Zhou B, Huang N, Farfel-Becker T, Sheng Z-H. Revisiting lamp1 as a marker for degradative autophagy-lysosomal organelles in the nervous system. Autophagy. 2018. Accessed August 5, 2023. https://www.ncbi.nlm.nih.gov/pmc/articles/PMC6103665/.

[51] Leica. Personal confocal promo. Personal Confocal Promo | Leica Microsystems. 2022. Accessed August 5, 2023. https://www.leica-microsystems.com/c/am/lsr-c/personal-confocal-promo/?nlc=20221205-SFDC-015774&utm_source=google&utm_medium=cpc&utm_campaign=23-AM-LSR-L3-STEL-GOOG-PP-STELLARIS-Search&utm_content=text_ad&utm_term=confocal+imaging&gclid=CjwKCAjwlJimBhAsEiwA1hrp5mwkfi_zp18sCtNir4oqillfxExUOglssfcImZqW0lX89qDychMs3hoChfEQAvD_BwE.

